# YibaoGaussian: Generate dynamic 3D models from text using Gaussian splatting as the explicit representation

**DOI:** 10.1101/2025.11.20.689534

**Authors:** Liu Mingtong, Huang Mengmeng, Olzhas N. Turar, Yang Chenghan, Ji Chaoqun, Xie Muhang

## Abstract

In recent years, the latest advances in the field of 3D content creation have been primarily based on neural radiation fields (NeRF). Despite significant research achievements, these methods often suffer from slow optimization speeds and high memory consumption. More importantly, these studies still mainly focus on static modeling, with very limited research on dynamic models, which restricts their practical applications. This paper proposes a method Yibaogaussian for generating dynamic 3D models from text descriptions, which can generate dynamic 3D models while ensuring efficiency and quality. Our key insight is on constructing canonical space and temporal deformation field . We build static 3D Gaussian fields in canonical space to capture the overall geometry and appearance features of a scene. By dynamically mapping the parameters of the Gaussian field to the temporal deformation field, we achieve continuous changes and natural motion of the static 3D model over time. Extensive experiments demonstrate that our proposed method offers higher efficiency and highly competitive generated quality. Notably, YibaoGaussian can generate high-quality dynamic 3D models from text descriptions in just 3 minutes, approximately 3 times faster than existing methods. This study verifies the feasibility of combining Gaussian representation and temporal deformation field in regular space, providing a new solution for text-driven dynamic 3D model generation and laying the technical foundation for multimodal applications such as digital humans, virtual reality (VR/AR), and film and television special effects.

## Introduction

With the rapid development of generative artificial intelligence [1], cross-modal generation from text-to-image [2] generation to 3D content modeling [3] has gradually become a research hotspot in the fields of computer vision and graphics. Diffusion models [4] have shown excellent performance in text-to-image generation tasks, enabling researchers to generate well-structured and realistic-looking 2D images based on natural language descriptions [5, 6].Inspired by this, a series of works such as DreamFusion and Magic3D combined the pre-trained text-to-image diffusion model with Neural Radiance Field (NeRF) to generate 3D models from text descriptions [7]. This type of method has achieved remarkable results in static scene modeling and provided a solid technical foundation for generating 3D content from text [8].At the same time, the explicit point representation method has shown great potential in the fields of static 3D reconstruction, real-time rendering, and especially in 3D Gaussian point rendering due to its efficient differentiable rasterization mechanism [9].However, existing research is mainly limited to static models and it is difficult to effectively describe dynamic three-dimensional models that change over time [10].For dynamic 3D models, how to ensure spatial geometric consistency [11] while maintaining smoothness in the time dimension [12] remains a core issue that needs to be solved. Traditional NeRF-based methods, such as D-NeRF [13] and HyperNeRF [14], have alleviated the temporal discontinuity phenomenon to a certain extent by introducing time conditions or deformation fields, but they still have problems such as slow optimization process and low rendering efficiency, which limit their feasibility in real-time interactive application scenarios. In addition, it is difficult to capture complex motion details by relying solely on implicit representation [15], while explicit representation [16] (such as three-dimensional Gaussian) is efficient but can not model temporal consistency, which can easily lead to jitter and loss of details in dynamic results [17]. To this end, this paper proposes a framework for text generation from dynamic 3D models based on Gaussian splatting representation. Specifically, a static 3D Gaussian representation is learned in a canonical space to capture the essential geometric and appearance features of the scene, and a temporal deformation field is introduced to simulate dynamic changes, generating a dynamic 3D model with temporal continuity and global consistency. At the same time, with the help of the semantic Destillation capability provided by the diffusion model, this study can effectively map natural language information into a dynamic three-dimensional space, thereby realizing the generation of high-fidelity models driven by text semantics. The contributions of this study are mainly reflected in two aspects: First, this study has achieved an important expansion of text-to-3D model generation from static scenes to dynamic content, providing new technical solutions for digital humans, virtual reality (VR/AR), film and television production, and interactive content generation;Secondly, the framework combines the implicit geometric modeling advantages of NeRF, the efficient differentiability of Gaussian rendering, and the semantic expression capabilities of the diffusion model, providing a feasible path for high-quality and efficient dynamic content generation. This study lays the theoretical and methodological foundation for future multimodal dynamic scene generation, such as speech-driven or action-driven dynamic 3D modeling.

## Related work

### Text to 3D Model

With the rapid development of diffusion models, research on text-driven 3D model generation has made significant progress [18].A typical method is DreamFusion [19], which uses the two-dimensional semantic prior provided by stable diffusion and score distillation sampling (SDS) technology [20] to refine text information into NeRF representation, thereby generating a 3D model from text descriptions. Subsequently, Magic3D [21] designed a two-stage optimization strategy: first, rapid convergence in low-resolution space, and then improving local details through high-resolution mesh optimization.Fantasia3D [22] combines the diffusion model with mesh-based optimization to achieve improved structural consistency and geometric accuracy.

Although these studies have advanced the progress of text-to-3D model generation, two limitations still exist: first, the generated results are mostly static models that lack temporal dimension modeling [23]; second, the implicit volume representation that NeRF relies on makes rendering expensive, which is not conducive to real-time or interactive applications [24].Recent work has begun to explore the direction of text-to-dynamic 3D models. For example, MAV3D [25] combines a diffusion model with temporal consistency constraints to achieve simple dynamic scene generation, but it is still limited in terms of geometric expression efficiency and scalability.Therefore, how to combine text-driven mechanisms with efficient dynamic modeling remains a core challenge that needs to be broken through [26].

### Dynamic scene modeling

Traditional NeRF models are mainly oriented towards static scenes and have limited ability to represent object motion or deformation [27].To solve this problem, D-NeRF [28] first introduced the concept of canonical space-time deformation field, decomposing the dynamic scene into a static reference configuration and a time-varying deformation field, thereby realizing the differentiable rendering of the model motion. This idea provides important inspiration for subsequent research.On this basis, T-NeRF directly uses time as an additional input dimension to explicitly represent the dynamic changes of the model, but due to the lack of a unified canonical space, it leads to poor geometric stability [29].HyperNeRF further models topological changes in high-dimensional latent space, enhancing its adaptability to complex motion and occlusion [30].DynIBaR uses video sequence feature interpolation to achieve dynamic reconstruction based on feature fields [31]. Although the above methods have made progress in temporal consistency and dynamic modeling, they still have obvious shortcomings: on the one hand, they lack the ability to drive text or semantics, making it difficult to control the generation of content [32]; on the other hand, the implicit representation based on NeRF has high computational overhead and slow rendering speed, making it unsuitable for real-time or large-scale generation tasks.

### 3D Gaussian Splatting

3D Gaussian Splatting, as an explicit point-based rendering method, provides a new solution for efficient 3D reconstruction and real-time rendering [33].Different from the implicit volumetric representation of traditional NeRF, our method decomposes the scene into a set of 3D Gaussian functions with position, covariance, and color properties, and enables fast and differentiable model generation via rasterization.Its main advantages are reflected in three aspects: (1) High efficiency: significantly improving training and inference speed and supporting real-time rendering; (2) Detail expression: able to capture high-frequency texture and edge features; (3) Scalability: able to naturally adapt to multi-view fusion and point cloud reconstruction tasks [34].Previous studies have shown that 3D Gaussian Splatting outperforms traditional methods in synthesizing new perspectives of static scenes [35]. However, this method still has shortcomings in dynamic modeling. Some works [36] attempt to introduce time-related parameters (such as position, covariance or color) for each Gaussian point to achieve time-varying rendering, but lack unified spatiotemporal constraints and global semantic consistency, which easily leads to artifacts, drift and temporal discontinuity in non-rigid scenes.

### Diffusion Model and Semantic Distillation

Diffusion models have become a core method in the field of image generation, which can generate semantically consistent and detailed two-dimensional images based on text input [37].To transfer this capability to 3D generation, the researchers proposed the SDS method: by optimizing the rendering results of the 3D representation to make it conform to the distribution of the diffusion model in the image domain, the mapping of text semantics to three-dimensional space can be achieved without 3D supervision.Subsequent research further improved the performance of SDS: (1)Variational fractional distillation (VSD) introduced probability distribution constraints to improve training stability [38]. (2)ControlNet+SDS fuses depth maps and edge information to enhance controllability [39]. (3)Latent Diffusion+3D scenes reduce the computational burden through latent space optimization. Nevertheless, SDS still faces two challenges: (1)The structural stability of the generated images is limited and it is easy to introduce ambiguity or fuzziness. (2)Its optimization is usually targeted at single-frame rendering, lacks temporal consistency constraints, and is difficult to directly apply to dynamic scene modeling.

### Problem Definition

The goal of this paper is to generate dynamic 3D models from text semantics. Formally speaking, given a piece of text semantics T, the model needs to learn a spatiotemporal mapping function,the function is as follows:

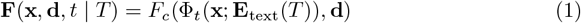

Here,x = (x, y, z) represents the position in three-dimensional space, d = (,) represents the line of sight of the camera, and t is the time variable.*E*_text_(T) encodes the semantics of the text,Φ_*t*_(**x**; *E*_text_(*T*)) is a time deformation field that maps a time point t to a canonical space. Compared with the task of generating static 3D models, generating dynamic 3D models from text has higher complexity and uncertainty.The main challenges are reflected in several aspects:First, due to the lack of direct 3D supervision signals for cross-modal semantic mapping, models usually need to rely on pre-trained diffusion models (e.g., Stable Diffusion) and use semantic distillation mechanisms to guide textual information into the 3D representation space.Second, non-rigid motion (e.g., human motion or cloth deformation) is common in dynamic models, which requires the ability to model continuous spatial deformation in the temporal dimension rather than relying solely on discrete frame interpolation. Third, directly regressing from five-dimensional input ((x, d, t)) to color and density output (c,) often leads to temporal redundancy and geometric instability problems. To this end, this study introduces a canonical space as a static benchmark and maps the model at any moment back to this unified space through a time deformation field, thereby ensuring spatiotemporal consistency and geometric continuity. Finally, traditional NeRF relies on implicit volume integration for rendering, which is computationally intensive and cannot meet the dual requirements of real-time and computational efficiency for dynamic generation tasks. To solve the above problems, this paper proposes a text-driven dynamic 3D model framework. Specifically, Stable Diffusion provides semantic prior guidance, while the NeRF module generates coarse geometric depth and occlusion information. Each 3D Gaussian point is then projected onto the screen plane to form a 2D elliptical splat. Depth sorting and alpha blending are then used to achieve front-to-back overlay and image synthesis. This framework effectively avoids volume integral calculations, significantly improves rendering efficiency, and enhances high-frequency detail performance while maintaining global low-frequency geometry. The canonical space at time t=0 is defined as a static benchmark model to maintain the consistency of the structure over time, the time deformation field learns the displacement mapping function from any time to the canonical space to capture the dynamic changes of non-rigid bodies.The overall mapping can be expressed as:

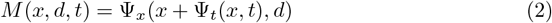

Here,Ψ_*x*_ predicts color and volume density, Ψ_*t*_ ensures dynamic consistency across time.

## Method

This chapter introduces in detail the overall structure and key modules of the proposed text-to-dynamic 3D model generation framework, The framework description is shown in Figure 1.The framework consists of five main components: (1) a semantic distillation module; (2) a NeRF backbone network in canonical space to capture low-frequency global geometric structure; (3) a 3D Gaussian layer in canonical space to enhance high-frequency details; (4) a temporal deformation field to simulate time-varying non-rigid motion; and (5) a dynamic rendering module that achieves efficient temporally consistent rendering through screen-domain rasterization. The framework description is shown in Figure 1.

**Fig 1.**
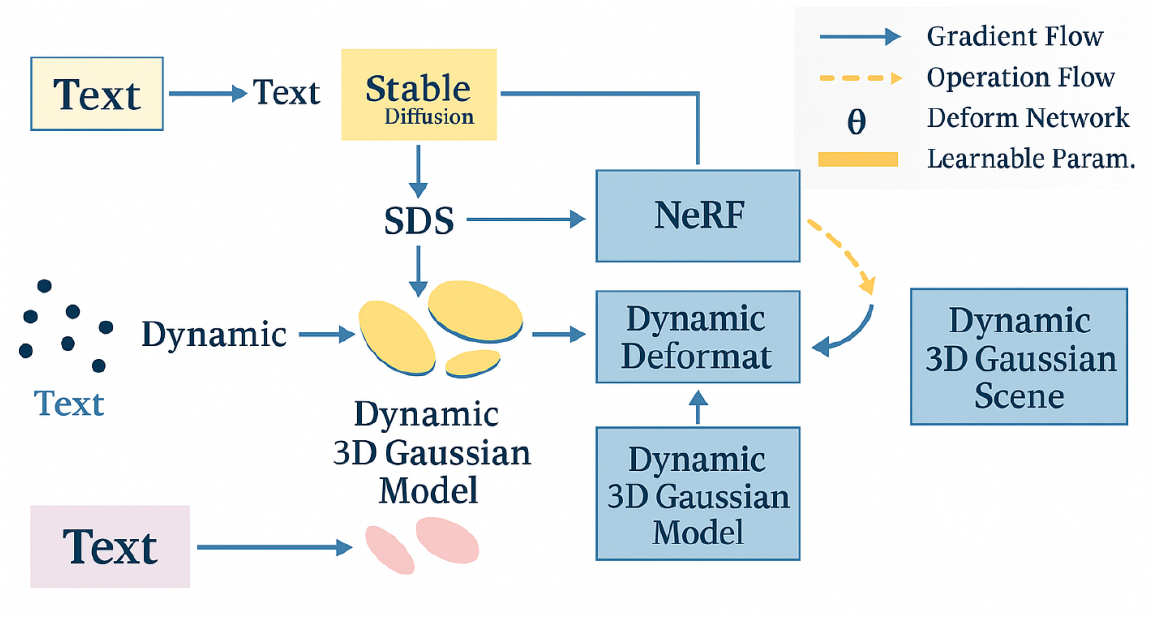
YibaoGaussian framework. The input text is processed using Stable Diffusion to extract cross-modal semantic priors, thereby obtaining geometric and appearance constraints consistent with the text description. This dynamic Gaussian representation is fed into NeRF and a dynamic deformation network, respectively. NeRF learns the implicit radiation field of the scene to supplement high-frequency structures, while the deformation network predicts the time-varying offsets of Gaussian points, thus capturing the dynamic changes of non-rigid bodies. The appearance structure from NeRF and the temporal offsets of the deformation field work together to form a 3D Gaussian representation, resulting in a dynamic 3D Gaussian scene with semantic consistency, geometric stability, and temporal continuity.

Figure 1.This paper organically combines the stable diffusion cross-modal semantic prior, the NeRF backbone network improved based on Gaussian representation, canonical space modeling and temporal deformation field to achieve the generation of dynamic three-dimensional models with temporal continuity from natural language descriptions. Text semantics are distillated into 3D representations through SDS, providing semantic guidance for NeRF and 3D Gaussian modules. The canonical space is used to capture the global structure of the static scene, and the temporal deformation field maps points at arbitrary time back to the canonical configuration to maintain geometric consistency. Dynamic Gaussian points are projected into screen space via differentiable Gaussian rasterization to generate high-fidelity dynamic images under adaptive density control. Finally, end-to-end optimization is performed through the joint loss function to achieve the generation of dynamic 3D models with consistent semantics, stable geometry and efficient rendering.

### Text semantic distillation module

In this study, the mapping of text semantics to 3D geometry is provided by a pre-trained stable diffusion model providing a semantic prior. we employ the SDS method to extract the semantic latent space of the diffusion model into a 3D representation. This process achieves knowledge transfer from text semantics to the 3D semantic space by optimizing the distribution consistency between the 3D model rendering and the image generated by the diffusion model. Specifically,For a given text T, a stable diffusion U-Net module is used to predict the noise and minimize the distribution difference between it and the rendered image latent variable,The diffusion model function is as follows:

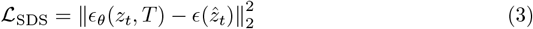

Here, *T* is the input text, *z*_*t*_ is the latent spatial noise of the diffusion model, *ϵ*_*θ*_ is the prediction noise function of stable diffusion, and 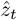 is the latent spatial variable of the rendering result. In order to enhance the stability of gradient propagation, this study introduces Variational Score Distillation,As shown in Formula 4:

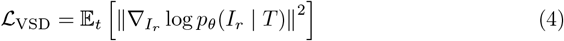

VSD effectively suppresses the gradient oscillation caused by blurred image samples by introducing variable score constraints in the latent space, thereby improving the robustness of semantic learning, stabilizing gradient propagation, and reducing the disturbance brought by blurred images.

### NeRF module

This paper uses the NeRF model to simulate the underlying geometry and illumination distribution. In addition, we define a static reference field in the gauge space to eliminate the temporal redundancy problem caused by dynamic deformation [35], instead of directly generating the dynamic model from NeRF. The input coordinates (x, d, t) are first mapped to the gauge space coordinates xc through the time deformation field, and the time deformation field function is as follows:

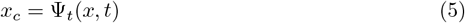

Where Ψ_*t*_ is the time deformation function The mapped coordinates are then fed into the NeRF network:

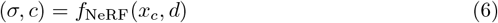

Where x represents volume density and y represents color. The rendering equation of NeRF is:

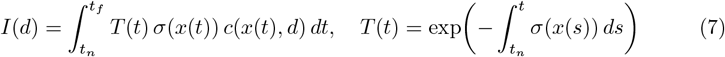

This module provides stable low-frequency geometry and overall lighting information, ensuring global consistency across different timeframes. At the same time, the introduction of canonical space effectively reduces the interference of model changes on network convergence, making time series modeling smoother. 4.3Gaussian explicit representation module (high-frequency details) To compensate for the shortcomings of NeRF in high-frequency texture and local details, we introduce a 3D Gaussian Splatting module in the canonical space as an explicit high-frequency detail layer. Each Gaussian point is represented by a triplet x, where y is the center position, z is the covariance matrix, and q is the color vector. By Gaussian rasterization, local texture, edge details, and surface reflectance characteristics can be explicitly modeled. he Gaussian distribution is defined as:

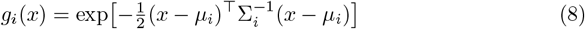

The final rendered color is a weighted superposition of all Gaussian contributions,The function is as follows:

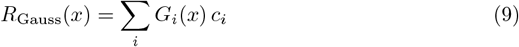

By explicitly modeling the point distribution, the network is able to effectively recover details, textures, and reflectance features.

### Time Deformation Field Module

To express the dynamic process that changes over time, we introduce a time deformation field Ψ_*t*_ to model the non-rigid displacement mapping of the scene from any time t to the canonical space.Its expression is:

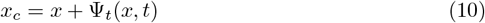

Here, x predicts the displacement vector for each spatial point in the time dimension.Adding a smoothness constraint x during training can ensure the temporal continuity and spatial consistency of the deformation.

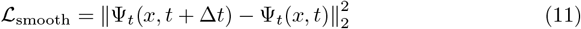

This mapping method from time to canonical space avoids the redundancy problem of directly learning the volume field in the time-deformation field while ensuring geometric consistency across time. It is a key module for realizing dynamic modeling.

### Dynamic rendering module

After completing semantic distillation (stable diffusion/SDS), geometric modeling (canonical NeRF), and deformation modeling (temporal deformation field), The goal of the rendering stage is to efficiently output the image results of each frame while ensuring temporal continuity and visual realism. To this end, we adopt Screen-Space Splatting as the core strategy to hierarchically fuse the volume rendering results of NeRF with the differentiable rasterization results of the Gaussian layer, while introducing temporal consistency and regularization terms to suppress cross-frame jitter and drift, thereby forming an end-to-end trainable dynamic rendering path. The method is divided into three parts according to the segmented structure: “screen rasterization multi-layer result fusiontemporal consistency and regularization”. The entire process consists of the following three stages.

#### Screen-Space Splatting

The spatial point at the current moment is mapped back to the canonical space through the time deformation field, and then converted into screen coordinates using the camera projection model.Each Gaussian point is rasterized into a 2D Gaussian kernel on the image plane in a differentiable manner, and pixel synthesis is achieved by combining depth sorting and alpha synthesis.This process effectively captures the relationship between surface reflectance and occlusion and enables high-quality visualization under the condition that gradients can propagate.In addition, in order to reduce high-frequency jitter and edge flickering, density adaptive control and visibility screening were introduced in the rendering process which significantly improved rendering efficiency and timing stability while ensuring details,Screen Space Rendering:

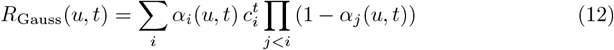

Here, 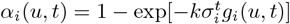 represents the transparency of the *i*-th Gaussian function at pixel *u*, 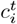 is the color of that Gaussian function, and it is sorted by depth.

### Gaussian display fused with NeRF output

NeRF is used to simulate basic geometry and lighting distribution, while the Gaussian module is responsible for simulating high-frequency details and edge textures.Model complementarity is achieved through weighted fusion, with the fusion weights adjusted according to the complexity of the scenario and the adaptive depth.The fused structure is shown in the figure 2.Meanwhile, the brightness and hue of the features are normalized before networking and fusion to ensure consistency of lighting and color, and to ensure that the output maintains visual continuity and stylistic consistency in the multi-layer structure,thereby,achieving a balance between efficiency and quality in generative models.

**Fig 2.**
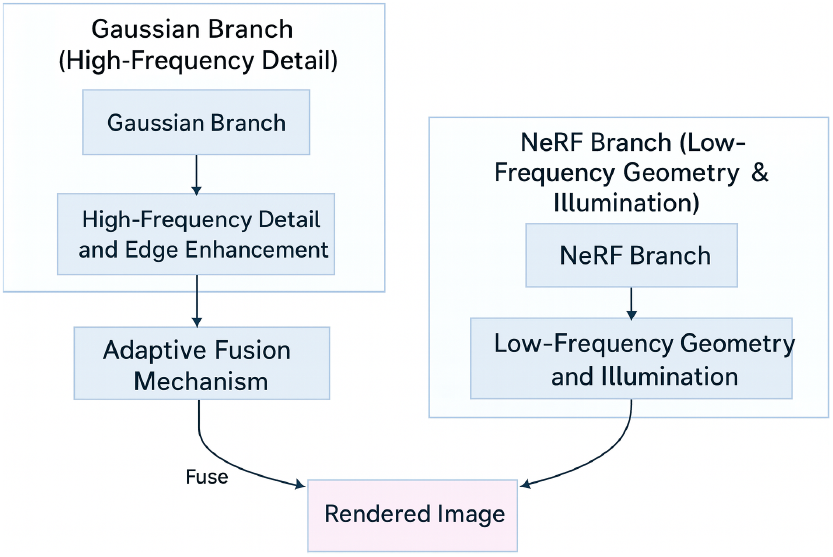
Gaussian display fused with NeRF output framework

**Fig 3.**
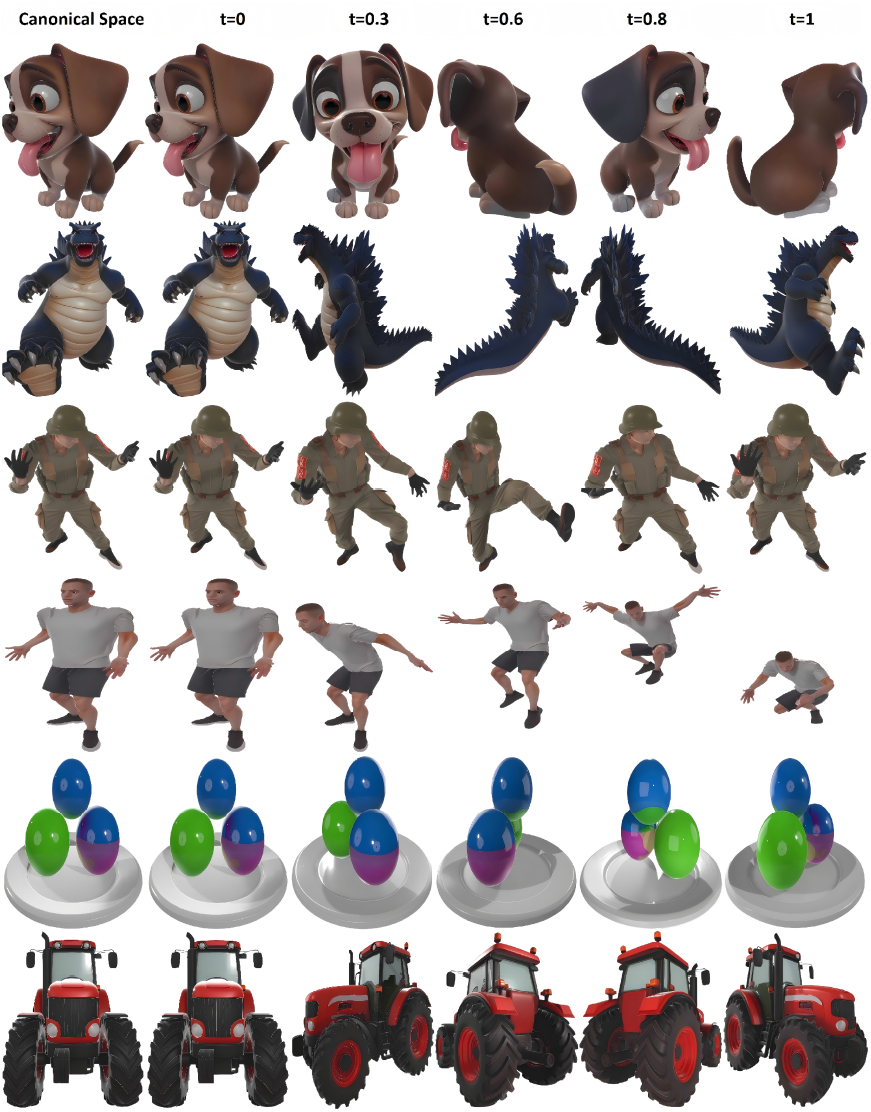
Samples generated by YibaoGaussian. Rows represent temporal changes, columns represent viewpoint changes, and the first column is the model in canonical space.

### Temporal consistency and regularization

The lack of constraints can easily lead to flickering and drifting between consecutive frames due to deformation and occlusion changes.To this end, this section introduces a temporal consistency loss term to constrain the smooth changes of adjacent frames in the rendered output.The network aligns the features of the previous frame with the current frame through the deformation field and minimizes the alignment error to maintain the continuity of motion and geometric structure.At the same time, a deformation smoothing regularization term is introduced to impose constraints on temporal and spatial gradients, making the transition of dynamic deformation more natural.Finally, semantic distillation loss, reconstruction error, temporal consistency, and deformation smoothness are integrated to form an overall optimization objective function to achieve end-to-end high-quality dynamic 3D model generation.In order to ensure the continuity of rendering results across frames, temporal consistency loss is introduced

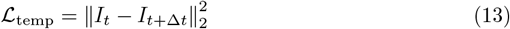

Align rendering results with real-world frame constraints

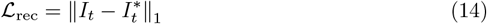

Combine semantic distillation, reconstruction error, temporal consistency, and deformation smoothness into an overall objective function

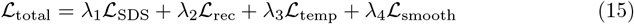

Here,x is the weight coefficient of each item, s is semantic consistency, d is graph reconstruction, f is temporal continuity, and g is deformation smoothness.

## Experiment

### Experimental Preparation

This study is implemented based on the PyTorch framework, and on this basis, the differentiable Gaussian rasterization module is improved to support time deformation fields and multi-stage visualization. The entire training process involves 40k iterations, of which the first 3k iterations only optimize the static 3D Gaussian field to obtain a stable spatial distribution and geometric shape. The 3D Gaussian field and the temporal deformation field are then jointly trained to achieve temporally consistent dynamic modeling. The optimizer uses a single ADAM with momentum parameters set to 1=0.9 and 2=0.999. Different modules use layered learning rates: 3D Gaussian splatting is consistent with the official implementation, The learning rate of the deformable network gradually decays exponentially from 8×104 to 1.6×106.During training, we jointly optimize semantic distillation and rendering losses and enable adaptive Gaussian density control to improve representation accuracy. Our study uses CLIP-O and CLIP-F [40] to measure the semantic relevance between generated content and textual prompts. We evaluate temporal consistency using the FVD [41] score to assess the quality and smoothness of rendered sequences. For qualitative analysis, we provided a visual comparative analysis and conducted a user study with 100 users. Users independently evaluate anonymized dynamic 3D model sequences generated by different models, rating them based on five attributes: 3D geometric consistency (3DC), appearance quality (AQ), motion fidelity (MF), text alignment (TA), and overall performance (OA) [42], and report the results as statistics for the Kicking Soldier model.

### Experimental implementation

To verify the effectiveness and robustness of the proposed YibaoGaussian framework in dynamic 3D content generation, we conducted experiments on a dynamic scene dataset provided by Diffusion4D, including rigid motion, non-rigid motion, and complex motion.The text prompts are described in English natural language. All images obtained by semantic distillation are downsampled to 400×400. 4096 rays are sampled in each iteration, and 128 points are uniformly sampled in each ray.The optimizer used was Adam, with an initial learning rate of 5×105 and an initial Gaussian number of 10K points. During training, the residual was used to trigger densification and splitting, and the model eventually converged to approximately 50K points.The experiments were conducted on an RTX 4090 GPU, and the training time for each model was approximately 3 days.

### Experimental results

#### Quantitative results

We compared four mainstream dynamic 3D models: D-NeRF, 3D-GS, DreamFusion, and Ours, and evaluated six dynamic scenes: Dog, StandUp, Godzilla, Tractor, Jumping, Hook, Monster, and BouncingBalls; and used three mainstream indicators: CLIP-F, CLIP-D, and FVD,The experimental results are shown in Table 1. On the CLIP-F and CLIP-O semantic consistency metrics, Ours achieved an average improvement of 4.3% and 6.7% over the second-place DreamFusion. On the FVD temporal quality metric, Ours achieved a 3.5% improvement over DreamFusion.Experimental results show that our model outperforms other models in terms of semantic alignment and temporal smoothness.In the generated Dog/StandUp, our results significantly outperform other methods on both CLIP-F and CLIP-O data, indicating that our method provides a more stable semantic mapping.In the generated Godzilla/tractor, DreamFusion exhibits a slight semantic drift.In the generated jumps/hooks, our method shows a significant reduction in FVD, indicating that our method is more stable in terms of temporal consistency and deformation modeling.In the generated monsters/bouncing balls, 3D-GS has stronger details but exhibits noticeable temporal jitter, while our model maintains the lowest FVD and the best smoothness.The temporal deformation field effectively simulates the continuity of motion and reduces frame jitter, especially in high dynamic models such as jumping and kicking.These results indicate that our YiBaoGaussian model achieves more consistent and realistic dynamic 3D reconstruction.

**Table 1.** Quantitative experimental results.

| Method | Dog |  |  | Godzilla |  |  | Kick |  |  |
| --- | --- | --- | --- | --- | --- | --- | --- | --- | --- |
|  | CLIP-F | CLIP-O | FVD | CLIP-F | CLIP-O | FVD | CLIP-F | CLIP-O | FVD |
|  | ↑ | ↑ | ↓ | ↑ | ↑ | ↓ | ↑ | ↑ | ↓ |
| D-NeRF | 0.735 | 0.568 | 497 | 0.708 | 0.568 | 537 | 0.700 | 0.562 | 503 |
| 3D-GS | 0.748 | 0.597 | 579 | 0.724 | 0.597 | 526 | 0.740 | 0.607 | 537 |
| DreamFusion | 0.807 | 0.624 | 482 | 0.793 | 0.622 | 473 | 0.805 | 0.634 | 475 |
| Ours | **0.834** | **0.672** | **465** | **0.825** | **0.697** | **445** | **0.837** | **0.663** | **457** |

| Method | Jumping |  |  | BouncingBalls |  |  | Tractor |  |  |
| --- | --- | --- | --- | --- | --- | --- | --- | --- | --- |
|  | CLIP-F | CLIP-O | FVD | CLIP-F | CLIP-O | FVD | CLIP-F | CLIP-O | FVD |
|  | ↑ | ↑ | ↓ | ↑ | ↑ | ↓ | ↑ | ↑ | ↓ |
| D-NeRF | 0.732 | 0.582 | 547 | 0.706 | 0.577 | 548 | 0.742 | 0.573 | 535 |
| 3D-GS | 0.769 | 0.637 | 534 | 0.737 | 0.606 | 573 | 0.774 | 0.607 | 572 |
| DreamFusion | 0.811 | 0.654 | 470 | 0.784 | 0.647 | 491 | 0.785 | 0.652 | 482 |
| Ours | **0.845** | **0.693** | **453** | **0.829** | **0.685** | **476** | **0.819** | **0.701** | **471** |

### Qualitative analysis

For a more intuitive evaluation, we provide qualitative results in the figure 2.These visual dynamics models highlight the ability of our approach to provide high-fidelity dynamic scene modeling.The figure depicts five scenes, drawn from several different perspectives at different times.The first column shows the model in canonical space at time t=0.Note that we are able to handle several types of dynamics: human motion in the jumping athlete and kicking soldier scenes; and the asynchronous motion of several bouncing balls.It is obvious from the results that our approach ensures enhanced consistency and captures complex rendering details in the new view rendering.

### User Research

We conducted a user study involving 100 participants and found that people showed stronger preferences for our approach across different attributes.As shown in Figure 4(a-c), in terms of 3DC, AQ, and MF, more users support YibaoGaussian, proving its ability to generate dynamic 3D models with realistic appearance, detailed geometry, and smooth motion.Experimental results show that the proposed method, YibaoGaussian, outperforms other methods on all metrics. The improvements in 3DC and AQ metrics are most significant, reaching 47% and 48% respectively, indicating progress in geometric reconstruction and appearance realism. On MF and TA metrics, our method achieves 47% and 46% respectively, demonstrating the effectiveness of temporal deformation fields in creating dynamic 3D models.YibaoGaussian achieves an all-round improvement from semantic consistency, geometric accuracy to dynamic timing by integrating NeRF’s low-frequency structure modeling, 3D Gaussian’s high-frequency detail enhancement and VSD semantic distillation, and the time deformation field alignment mechanism, demonstrating the significant advantages of the framework proposed in this paper in dynamic 3D generation tasks.

**Fig 4.**
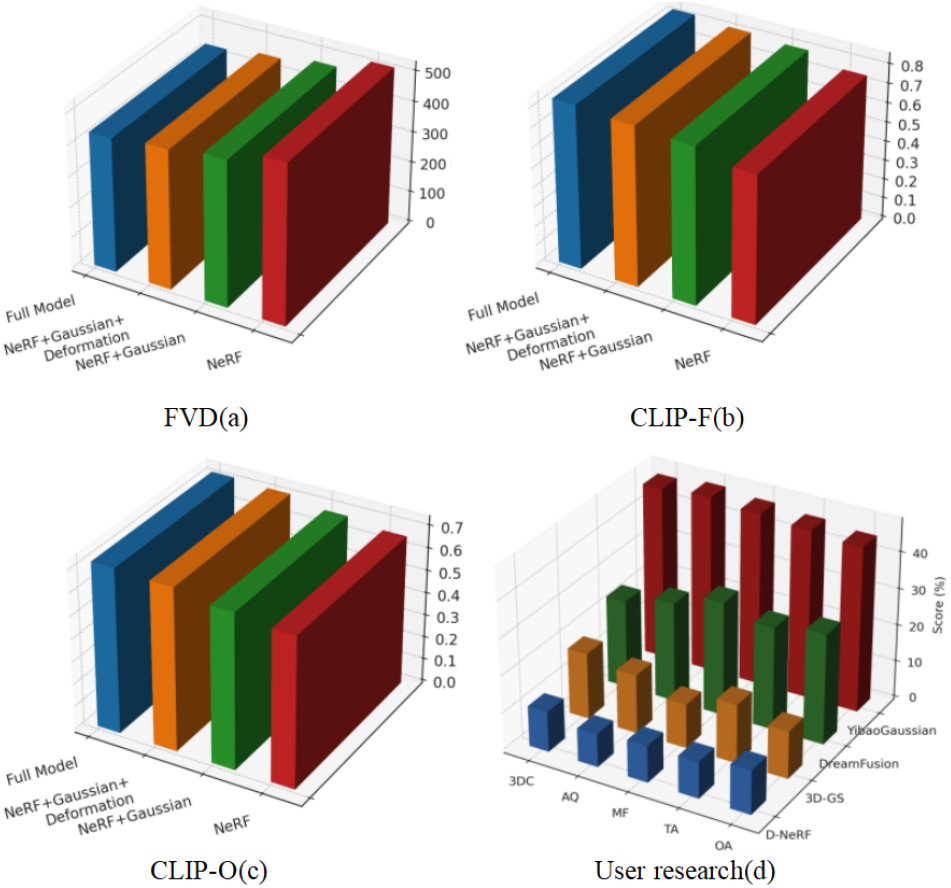
(a)(b)(c)Ablation experiment results. The results demonstrate a comparison of different component combinations across three metrics for 3D model performance. CLIP-F and CLIP-O reflect the model’s performance in terms of semantic consistency and semantic controllability. In the FVD metric, which measures temporal consistency, the complete model also achieved the lowest score, further validating the effectiveness of the proposed framework in terms of continuity and stability in dynamic 3D generation tasks.(d)User survey results. Distribution of preference ratings generated by the same model from 100 users using different methods.

### Ablation experiments

To analyze the impact of each module on overall performance, we conducted ablation experiments at different training stages.The results of the jumping man model under different modules are shown in Figure 4(d), and the visualization comparison is shown in Figure 5.(1)ithout screen domain rasterization. The number of training iterations was the same as YibaoGaussian. It can be seen that, in terms of text alignment, model quality, and motion effects, Jumping Boys tends to use the screen domain rasterization training method.(2)Without time deformation field. The absence of a time-deformation field will lead to a slower convergence rate and increase dynamic instability.(3) Without Gaussian. The lack of Gaussian will significantly reduce high-frequency details and texture fidelity, resulting in a very blurry model.Our findings indicate that the omission of any design module leads to a decrease in the quality of the generated model.

**Fig 5.**
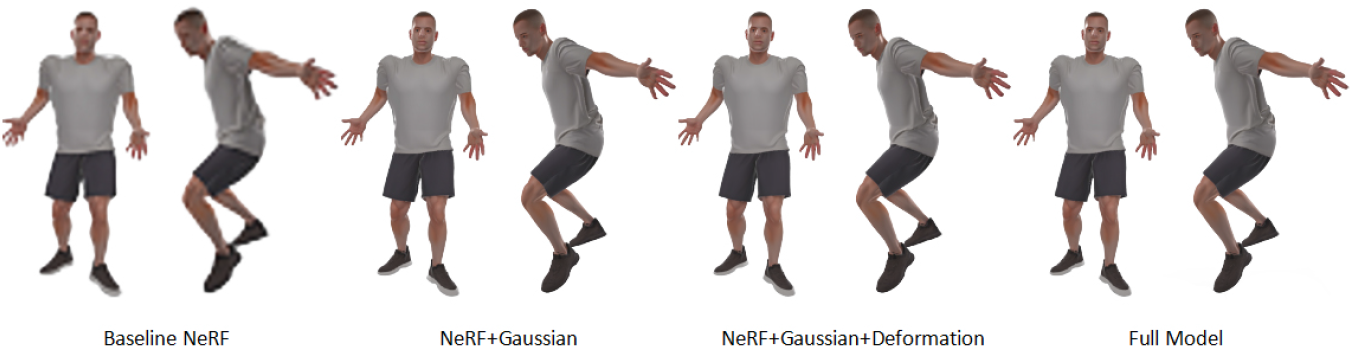
Visual comparison of ablation experiments

Specifically, the final dynamic 3D model exhibits greater ambiguity and inaccuracy.

We verify the effectiveness of our model by conducting systematic experiments on various synthetic and real dynamic models.Our framework not only outperforms existing methods in image quality and semantic alignment, but also achieves significant improvements in dynamic consistency and rendering efficiency, providing a new solution for text-driven dynamic 3D content generation that balances accuracy and efficiency.Experimental results verified in multiple scenarios and indicators fully demonstrate the superiority of the proposed YibaoGaussian framework.The framework not only surpasses existing models in semantic consistency and rendering quality, but also strikes a balance between dynamic continuity and real-time performance.In practical applications, YibaoGaussian can be widely used in digital human modeling, virtual scene generation, interactive content production, etc., providing an efficient and scalable technical solution for dynamic 3D model generation.

## Conclusion

This paper proposes YibaoGaussian, a unified framework for text-to-dynamic 3D model generation.This method organically combines the stable diffusion cross-modal semantic prior, the improved NeRF based on Gaussian representation, the canonical space and the temporal deformation field, and realizes the direct generation of dynamic three-dimensional models with temporal continuity from natural language descriptions. We systematically validated our method using real-world datasets. Experimental results demonstrate that our method outperforms existing methods in terms of model quality, semantic consistency, temporal stability, and rendering efficiency. The collaborative work of multiple modules enables our method to achieve a good balance between textual semantics, geometric structure, and rendering efficiency. Our research not only improves the quality and efficiency of existing text-to-dynamic 3D model generation tasks, but also provides a scalable solution for text-driven dynamic 3D model generation. Our core innovation is the proposal of a “moving query instead of moving representation” mechanism, which effectively improves consistency in the time dimension and, combined with explicit Gaussian rendering, ensures the uniformity of dynamic model quality and structure. Our approach is applicable not only to simple human and animal movements, but also inspires future complex large-scale interactive scenarios and large-scale video generation tasks. Looking ahead, YibaoGaussian still has room for expansion: (1) Introducing physical constraints into gauge space to enhance the physical properties of complex dynamics.(2) Explore deep integration with video generation models to achieve joint optimization from text to dynamic models to video.(3) Further improve the scalability of Gaussian rendering so that the model can run efficiently on mobile devices or real-time virtual reality. In summary, our approach advances the technology of text-to-dynamic 3D model generation, provides a new solution for multimodal dynamic 3D modeling, and lays a solid foundation for building next-generation intelligent interactive 3D content generation systems.

## Notes

### Competing Interest Statement

The authors have declared no competing interest.

